# Regulation Network of Colorectal Cancer Specific Enhancers in Progression of Colorectal Cancer

**DOI:** 10.1101/2021.06.14.448310

**Authors:** Bohan Chen, Yiping Ma, Jinfang Bi, Wenbin Wang, Anshun He, Guangsong Su, Zhongfang Zhao, Jiandang Shi, Lei Zhang

**Affiliations:** State Key Laboratory of Medicinal Chemical Biology and College of Life Sciences, Nankai University, 94 Weijin Road, 300071 Tianjin, China

**Keywords:** Colorectal cancer-specific enhancer, long-range interaction, Hi-C, HiChIP, TAD, loop

## Abstract

Enhancers regulate multiple genes through higher-order chromatin structure and further affect cancer progression. Epigenetic changes in cancer cells activate several cancer specific enhancers that are silenced in normal cells. These cancer specific enhancers are potential therapeutic targets of cancer. However, functions and regulation network of colorectal cancer specific enhancers are still unknown. Here in this study, we profile colorectal cancer specific enhancers and reveal the regulation network of these enhancers by analysis of HiChIP, Hi-C and RNA-seq data. We propose the regulation network of colorectal cancer specific enhancers plays important role in progression of colorectal cancer.

## 1. Introduction

Colorectal cancer is the world’s fourth most deadly cancer [1]. Previous studies on colorectal cancer indicated sequential accumulation of genetic mutations and chromosomal instability are the important reasons for the occurrence and development of colorectal cancer [2]. Typical genomic events in the initiation of colorectal cancer are *APC* mutation and followed by *RAS* activation or function loss of *TP53* [1]. Besides, *MAPK*, *WNT*, *PI3K*, TGF-βsignaling pathway are reported play important roles in initiation and progression of colorectal cancer [2, 3]. Chromosomal changes including deletion, amplifications and translocations induced alterations of genes’ expressions and functions that could further initiate colorectal cancer. Studies on higher-order chromatin structures revealed long-range gene regulation networks are important in modulating biological process in cells [4–6]. Further studies on gene regulation networks could help unveil mechanism of occurrence and progression of colorectal cancer.

Gene regulatory elements like enhancers or silencers could regulate multiple target genes through higher-order chromatin structures [7, 8] and function in gene regulation networks in cancer cells. Mutations in gene regulatory elements and changes of higher-order chromatin structures are related to gene transcriptions and cancer progression [8–10]. Prior study revealed reorganizations of chromatin structure in colorectal cancers compared with normal colons. And the topological changes of chromatins restrain malignant progression of colorectal cancers [11]. In addition, changes of higher-order chromatin structures in colorectal cancer cells are accompanied by epigenetic changes like alterations of DNA methylation [11, 12]. These epigenetic changes are associated with changes of transcription factors’ binding and activation of some gene regulatory elements like enhancers. Some enhancers are activated in cancer cells while not in normal cells [13]. These “cancer specific enhancers” may regulate several target genes through higher-order chromatin structures and play roles in cancer progression. These cancer specific enhancers are potential therapeutic targets of cancer. However, how colorectal cancer specific enhancers functions in cancer initiation and progression is still not clear.

In this study, by combination analysis of ChIP-seq, HiChIP and Hi-C data, we show profile of colorectal cancer specific enhancers and gene regulation network of these enhancers. Results in this study suggest that colorectal cancer specific enhancers could regulate multiple target genes including cancer driver genes, and genes regulated by these colorectal cancer specific enhancers are associated with occurrence and development of colorectal cancer.

## 2. Results

### 2.1 H3K27ac profiles define colorectal cancer specific enhancers

H3K27ac is a well-characterized histone modification marker of active enhancers[14]. To identify colorectal cancer-specific enhancers, we analyzed H3K27ac CHIP-seq datasets from 7 published colorectal datasets and 10 normal tissues from the NIH Roadmap database [15]. Unsupervised hierarchical clustering was performed on H3K27ac enrichment loci which identified 10 clusters from the 59 samples (Fig. 1A) and show distribution pattern of enhancers was highly tissue specific, which is consistent with previous studies [13]. 21132 normal colon tissue specific enhancers and 11463 colorectal cancer specific enhancers were identified. And the *t*-distributed stochastic neighbor embedding (*t*-SNE) analysis indicated colorectal cancer enhancer profiles were distinct from normal colon tissue enhancers (Fig. 1B). In addition, we identified 763 super enhancers among the colorectal cancer specific enhancers (Fig. 1C). Some of these super enhancers are associated with oncogenes, like *EPHA2*, *LIF*, *ID1*, *SMAD7*, *BMP4*, *FOXA1*, *NOTCH1*.

**Fig 1.**
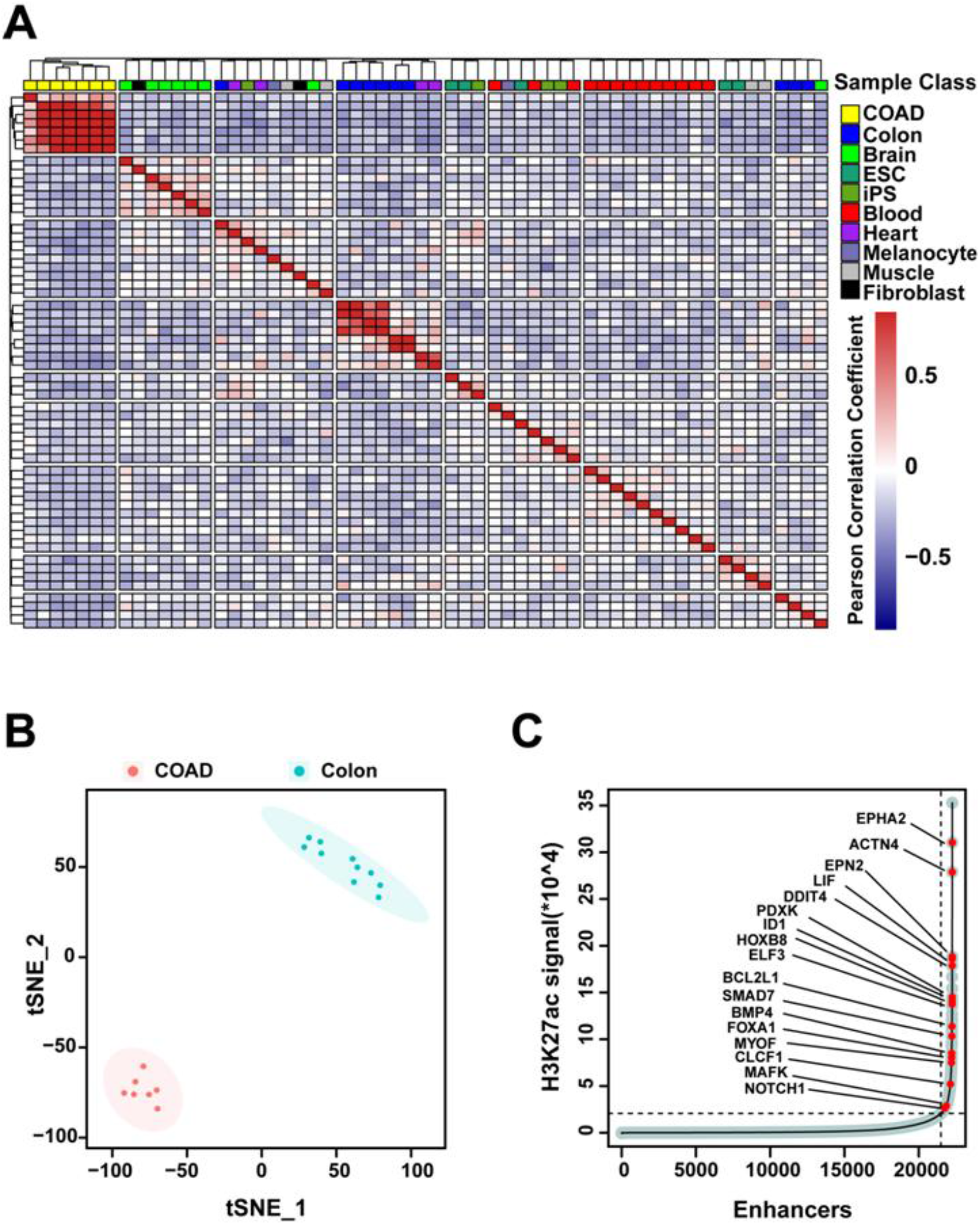
H3K27ac profiles define colorectal cancer specific enhancers. A) Unsupervised hierarchical clustering of the 11463 enhancer loci detected in colorectal cancer(n=7) compared to the normal tissue samples(n=10). B) *t*-distributed stochastic neighbour embedding (*t*-SNE) analysis of normal colon tissue specific enhancers and colorectal cancer specific enhancers. C) Inflection plot showing identified super enhancers among the cancer-specific enhancers.

### 2.2 HiChIP identifies chromatin interactions containing colorectal cancer specific enhancers

Positions of colorectal cancer specific enhancers are shown in Figure 2A, these colorectal cancer specific enhancers are evenly distributed on each chromosome (Fig. 2A). To identify the regulation network of colorectal cancer specific enhancers, HiChIP for H3K27ac was performed in colorectal cancer cell line HCT116. HiChIP data indicate interaction loops between enhancers/super enhancers and other chromatin fragments (Fig. 2B, C, interaction loops in chromosome 1 were shown as an example). And colorectal cancer specific enhancers/super enhancers also interact with several chromatin fragments including some long-range chromatin regions (Fig. 2D, E, interaction loops in chromosome 1 were shown as an example). Most of the interaction loops that contain colorectal cancer specific enhancers are less than 100kb in length and include more short-range interactions than random interaction loops from the HiChIP data (Fig. 2F). In addition, for the interaction loops that are more than 100kb in length, random longer interaction loops (>100kb) have a lower proportion (Fig. 2G), but longer interaction loops (>100kb) that contain colorectal cancer specific enhancers didn’t have a significant lower proportion (Fig. 2G), which suggests more long-range interactions are associated with colorectal cancer specific enhancers.

**Fig 2.**
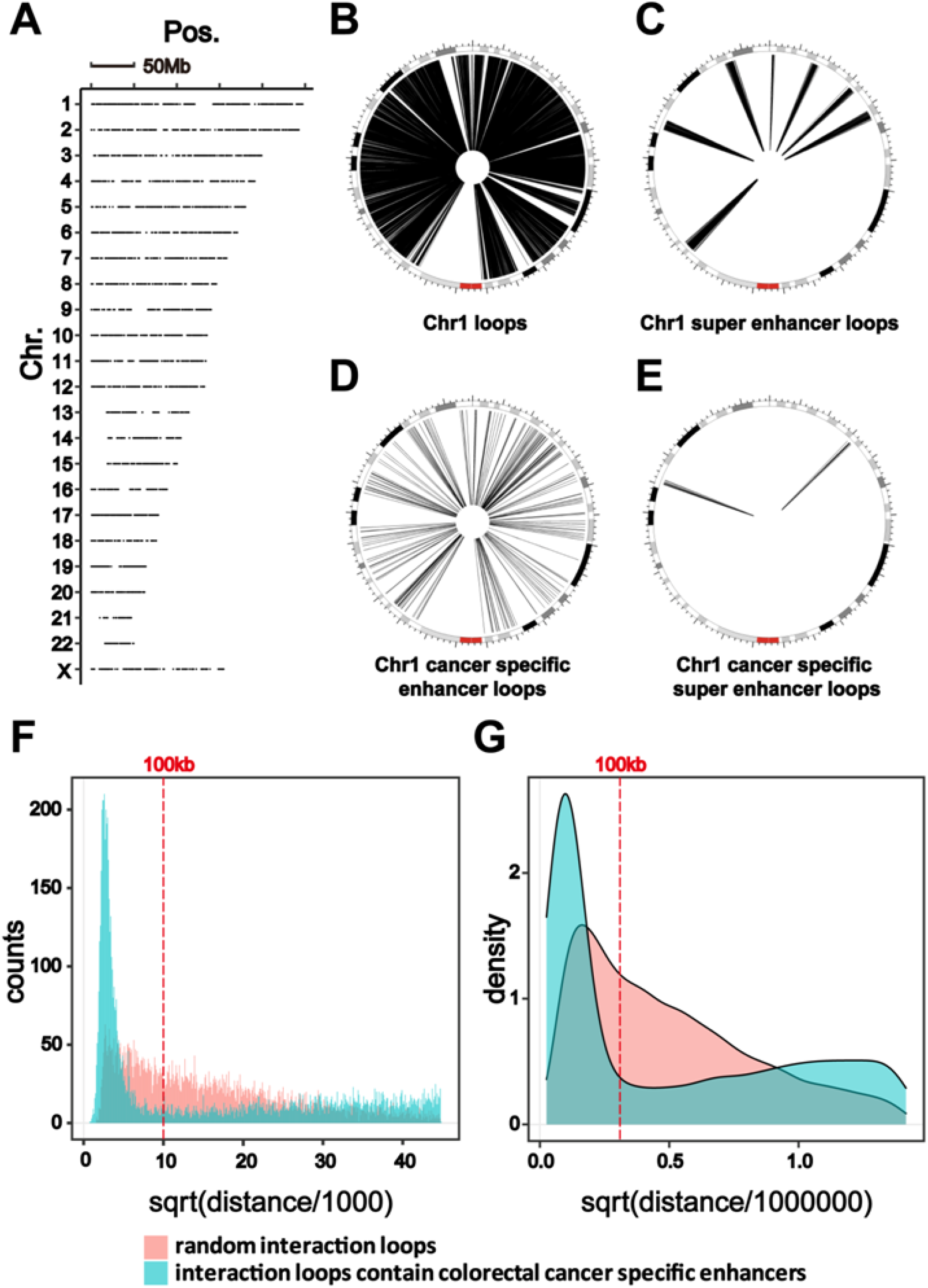
HiChIP identifies chromatin interactions containing colorectal cancer specific enhancers. A) Distributions of colorectal cancer specific enhancers on human genome (hg19). B and C) Circos plot showing interactions in Chr1 indicated by curves extending from enhancers and super enhancers in colorectal cancer cells. D and E) Circos plot showing interactions in Chr1 indicated by curves extending from colorectal cancer specific enhancers and super enhancers in colorectal cancer cells. F and G) Analysis of length distribution of random interaction loops from HiChIP data (Red) and interaction loops that contain colorectal cancer specific enhancers (Blue).

### 2.3 Change of TADs’ boundaries in colorectal cancer cells compared with normal colon cells

Based on analysis of Hi-C data in colorectal cancer cells and normal colon cells [11], about 65% boundaries of TADs changed in colorectal cancer cells compared with normal colon cells (Fig. 3A-F). The boundary-changed TADs were divided into six categories: expand (Fig. 3A), narrow (Fig. 3B), fusion (Fig. 3C), split (Fig. 3D), appear (Fig. 3E) and disappear (Fig. 3F), that respectively represent six kinds of change patterns of TADs’ boundaries in colorectal cancer cells compared with normal colon cells. Previous studies have indicated that gene transcription is correlated with insulation at TAD boundaries [6, 11]. Expand, fusion and appear of TADs in colorectal cancer cells are associated with more interactions between enhancers and genes, while split, narrow and disappear are associated with less interactions between some enhancers and genes in colorectal cancer cells. 65% of changed TADs in colorectal cancer cells compared with normal colon cells suggest global alterations of gene regulations in colorectal cancer cells. We noticed that more than 50% (51%-86%, except for the “disappear” group, which is 24%) of these changed TADs containing colorectal cancer specific enhancers (Fig. 3A-F), which suggests important roles of colorectal cancer specific enhancers in the alterations of the gene regulation network.

**Fig 3.**
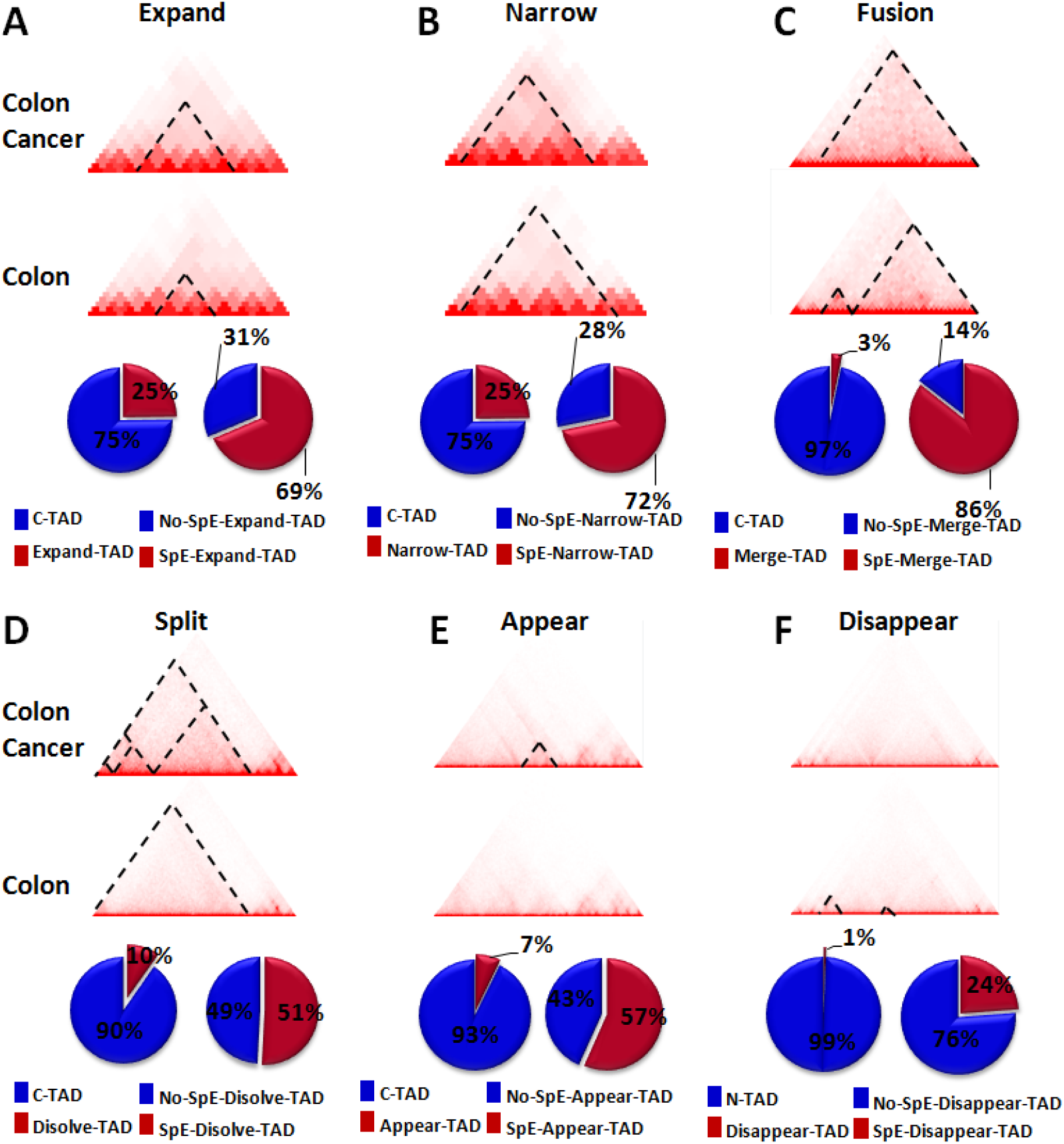
Change of TADs’ boundaries in colorectal cancer cells compared with normal colon cells. A) “Expanded TADs” in colorectal cancer cells compared with normal colon cells. B) “Narrowed TADs” in colorectal cancer cells compared with normal colon cells. C) “Fusion TADs” in colorectal cancer cells, two or more TADs in normal colon cells fused to one TAD in colorectal cancer cells. D) “Split TADs” in colorectal cancer cells, new insulation boundaries were formed in one TAD in colorectal cancer cells. E) “Appeared TADs” in colorectal cancer cells that are not exist in normal colon cells. F) “Disappeared TADs”, TADs in normal colon cells that are disappeared in colorectal cancer cells. Pie charts in A-F show the percentages of specific category of boundary-changed TADs (category-TAD) in total changed TADs (C-TAD) in colorectal cancer cells and the percentages of specific category of boundary-changed TADs contain colorectal specific enhancers (SpE-category-TAD) in total changed TADs of this category.

### 2.4 Transcriptome change in colorectal cancer cells is associated with colorectal specific enhancers

To figure out whether transcription change of genes in colorectal cancer cells were associated with colorectal specific enhancers, we analyzed the transcriptome data of colon tissues (n=349) and colorectal cancer tissues (n=275) from TCGA. 4714 genes were significantly upregulated (fold change>1.5, q<0.01) and 4924 genes were significantly downregulated (fold change>1.5, q<0.01) in colorectal cancer tissue (Fig. 4A). Combination analysis of RNA-seq and HiChIP data identified 152 genes are significant upregulated (fold change>1.5, q<0.01) in colorectal cancer tissue and have comparatively high interaction signal (interaction counts ≥5) with colorectal cancer specific enhancers (Fig. 4B). These genes are the potential target genes that are directly regulated by colorectal cancer specific enhancers. About 79% of the interactions between colorectal cancer specific enhancer and target genes are long-range (≥20kb) interactions (Fig. 4C). Pathway and process enrichment analysis by Megaspace [16] showed potential target genes of colorectal cancer specific enhancers are enriched in RNA catabolic process, α6β1α6β4 integrin pathway and regulation of cyclin-dependent protein serine/threonine kinase activity (Fig. 4D). These pathways or biological processes are reported associated with colorectal cancer progression [17–21]. *ITGB4* encodes integrin subunitβ4 and was involved in the α6β1α6β4 integrin pathway.

**Fig 4.**
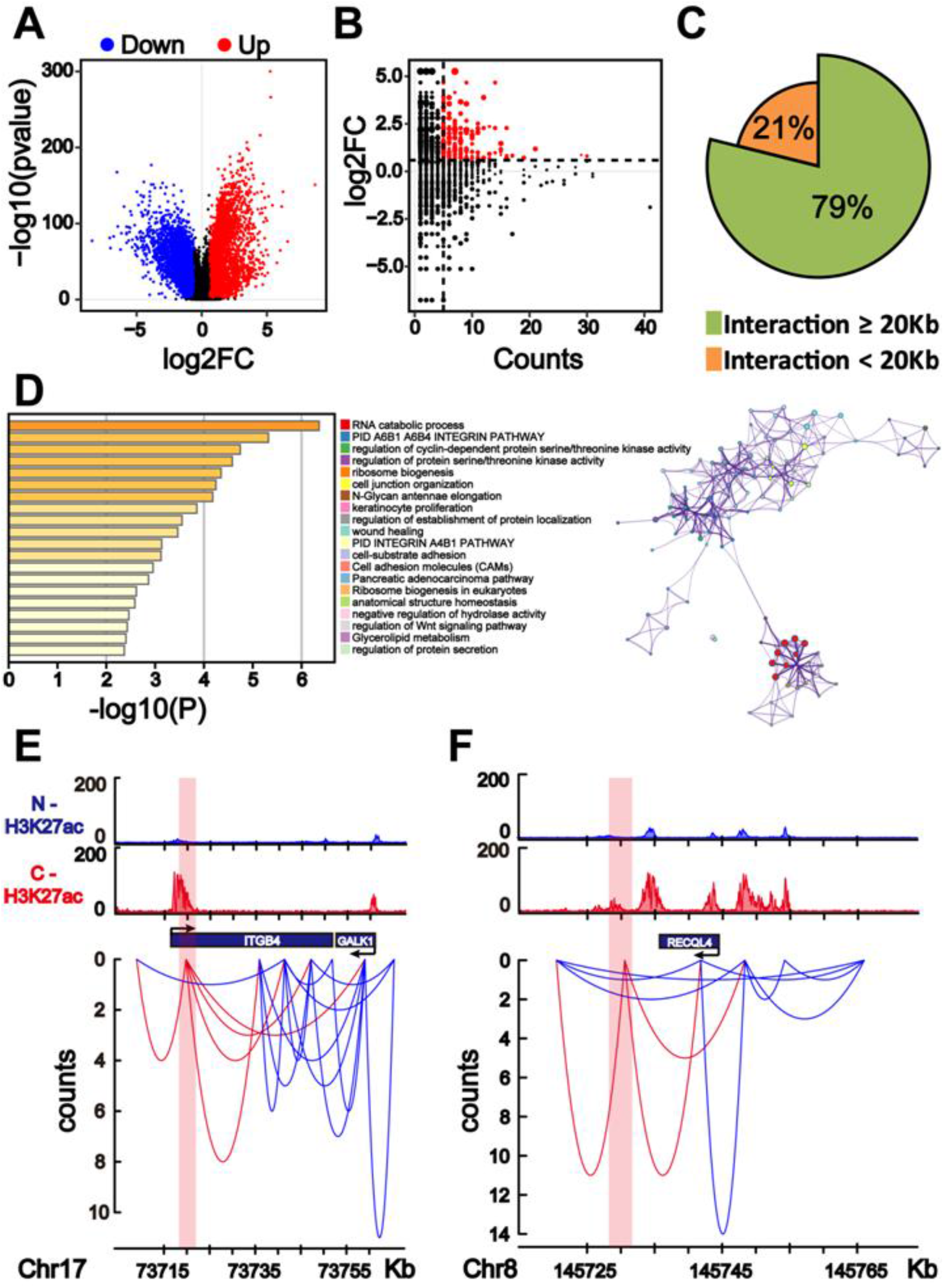
Transcriptome change in colorectal cancer cells is associated with colorectal specific enhancers. A) Volcano plot depicting gene expression changes in normal colon tissue (n=349) and colorectal cancer tissue (n=275). B) Comparison of fold-changes in gene expression based on RNA-seq and HiChIP signal. 735 target genes are shown on the plot. Red plots indicate target genes that significantly changed (fold change>1.5, q<0.01) in colorectal cancer tissue compared with normal colon tissue and have significant HiChIP signal (counts ≥ 5). C) Percentage of long-range (>20kb) interaction loops between colorectal cancer specific enhancers and target genes. D) Functions of genes as analyzed by Megaspace. E) H3K27ac enrichment for normal colon (N, blue) and colorectal cancer (C, red) and HiChIP interaction loops (red curves) between target gene *ITGB4* and colorectal cancer specific enhancer. Colorectal cancer specific enhancer is shaded in red. Blue curves indicate other interaction loops in this area. F) H3K27ac enrichment for normal colon (N, blue) and colorectal cancer (C, red) and HiChIP interaction loops (red curves) between target gene *RECQL4* and colorectal cancer specific enhancer. Colorectal cancer specific enhancer is shaded in red. Blue curves indicate other interaction loops in this area.

HiChIP data show *ITGB4* is regulated by a colorectal cancer specific enhancer (Fig. 4E) and *ITGB4* was upregulated in colorectal cancer tissue based on RNA-seq data, suggestive of regulation of *ITGB4* by colorectal cancer specific enhancer. Mutations and aberrant expression of cancer driver genes affect key cellular functions and induce cancer occurrence. *RECQL4* is one of the cancer driver genes [22] and potential target genes that are regulated by colorectal cancer specific enhancer (Fig. 4F). In addition, *RECQL4* is one of the most often mutated genes in colorectal cancer [23]. The colorectal cancer specific enhancer is about 15kb away from the *RECQL4* promoter and HiChIP data show interactions between colorectal cancer specific enhancer and *RECQL4* (Fig. 4F).

## 3. Discussion

This study shows colorectal specific enhancers interact with several potential target genes that are upregulated in colorectal cancers compared with normal colon. HiChIP data indicate higher-order chromatin structure induce interactions between colorectal cancer specific enhancers and target genes. Most altered TADs in colorectal cancer genome contain colorectal specific enhancers. But it is not clear whether changes of chromatin topological structures or the activation of colorectal specific enhancers is the primary event in the initiation of colorectal cancer. Previous study showed that chromosomal topological changes repress stemness and invasion programs and may restrain malignant progression of colorectal cancer and tumor-associated epigenomic changes are primarily oncogenic [11]. HiChIP data show that some of target genes of colorectal cancer specific enhancers are cancer driver genes, including *RECQL4* that is showed in Figure 4F, and analysis of Hi-C data indicate most of boundary-changed TADs contain colorectal specific enhancers (Fig. 3). It is possible that the activation of colorectal specific enhancers is before alteration of higher-order chromatin structure during occurrence and progression of colorectal cancer. Further studies are needed to verify this hypothesis.

With the development of chromatin conformation capture techniques [24–27], analysis of higher-order chromatin structures in cancer cells revealed complicated long-range gene networks of enhancers and indicated important functions of these enhancers or gene networks in cancer progression [8–10, 13]. A recent study reported that mutation in enhancer loci leads to activity change of the enhancer and affects expression of long-range target genes of this enhancer and further improve the progression of prostate cancer [8]. Prior study showed that ependymoma enhancer profiles were distinct from other tissues and most super enhancers were tumor-specific and enriched with cancer-associated genes [13], suggestive of important functions of these tumour-specific enhancers in the initiation of tumor. Our study profiles the colorectal cancer specific enhancers and presents regulation network of these enhancers based on HiChIP data and RNA-seq data, further unveils the mechanism of initiation and progression of colorectal cancer.

Combination analysis of HiChIP data and RNA-seq data indicated several potential target genes of colorectal specific enhancer. *ITGB4* is one of these potential target genes. ITGB4 is one of integrin (ITG) molecules that are associated with cell migration, proliferation and cancer development [28–30]. Aberrant *ITGB4* expression was reported in several cancers including colorectal cancers [18, 31]. RNA-seq analysis also showed significant upregulation of *ITGB4* in colorectal cancer tissue. A recent study indicated that ITGB4 might be a prognosis marker for individual therapy of colon cancer [18]. Data in our study suggests upregulation of *ITGB4* in colorectal cancer cells may due to the regulation of colorectal cancer specific enhancer. In addition, we noticed that *GALK1*, which is beside *ITGB4* (Fig. 4E), may also be one of the potential target genes of the colorectal cancer specific enhancer which is showed in Figure 4E. HiChIP data indicate interactions between the colorectal cancer specific enhancer and *GALK1*, and RNA-seq analysis showed upregulation (fold change=1.97, q=7.76×10^−49^) of *GALK1* in colorectal cancer tissue compared with normal colon tissue. However, the interaction signal is not very robust in the HiChIP data (count=3). This may due to the long-range interactions between the colorectal cancer specific enhancer and *GALK1* (~45kb). Our study suggests *GALK1* may be regulated by colorectal cancer specific enhancer and associated with progression of colorectal cancer. However, few studies reported function of *GALK1* in colorectal cancer for now. *RECQL4* is an example of cancer driver genes that are regulated by colorectal cancer specific enhancers. RECQL4 was reported as DNA helicase and function in DNA replication, DNA repair and recombination [32], which play important role in maintaining the genomic stability [33]. Several studies indicated RECQL4 is associated with cancer progression and could be diagnostic marker for cancer [33–37]. Examples include overexpression of *RECQL4* is associated with poor prognosis of gastric cancer [35] and predicts poor prognosis in hepatocellular carcinoma [36]. Investigation of mutation spectrum of cancer-associated genes in patients with early onset of colorectal cancer showed that *RECQL4* is one of the most often mutated genes [23]. And analysis of RNA-seq data from 275 colon cancer tissues and 349 normal colon tissues show that *RECQL4* is upregulated in colorectal cancer cells. Based on literatures and data analysis in the present study, it suggests *RECQL4* is an important target gene of colorectal cancer specific enhancers and involved in the progression of colorectal cancer. Although further studies are needed to pinpoint whether there are key driver genes in the colorectal cancer specific enhancer regulation network or the progression of colorectal cancer a combined result of these target genes, our study suggests important functions of colorectal cancer specific enhancers in colorectal cancer progression, and these enhancers could be potential therapeutic targets of colorectal cancer.

In summary, our study shows that colorectal cancer specific enhancers regulate several target genes including cancer driver genes and may play important roles in occurrence and progression of colorectal cancer.

## 4. Materials and methods

### 4.1 Cell Culture

The colorectal cancer cell line HCT116 was obtained from the ATCC. HCT116 cells were incubated at 37°C with 5% CO_2_ and cultured in Dulbecco’s Modified Eagle Medium (GIBCO) medium, supplemented with 10% fetal bovine serum (FBS) (BI) and 1% penicillin-streptomycin (GIBCO).

### 4.2 HiChIP

HiChIP was performed as described previously with modifications [26]. Briefly, cells were crosslinked with 1% formaldehyde and lysed. Then the chromatin was digested using *Mbo*I (NEB), and the restricted ends were religated by T4 ligase (NEB). Pelleted nuclei were dissolved in nuclear lysis buffer (50 mM Tris-HCl, pH 7.5, 10 mM EDTA, 1% SDS and protease inhibitors) and were sonicated and diluted in ChIP Dilution Buffer (0.01% SDS, 1.1% Triton X-100, 1.2 mM EDTA, 16.7 mM Tris-HCl, pH 7.5, and 167 mM NaCl). Then immunoprecipitation was performed overnight at 4 °C by incubating H3K27ac antibody (Abcam) precoated on protein A-coated magnetic beads (Thermo Fisher Scientific). Immunocomplexes were washed three times each with low-salt buffer (0.1% SDS, 1% Triton X-100, 2 mM EDTA, 20 mM Tris-HCl, pH 7.5 and 150 mM NaCl), high-salt buffer (0.1% SDS, 1% Triton X-100, 2 mM EDTA, 20 mM Tris-HCl, pH 7.5 and 500 mM NaCl), LiCl buffer (10 mM Tris-HCl, pH 7.5, 250 mM LiCl, 1% NP-40, 1% Na-Doc and 1 mM EDTA). Beads were resuspended in DNA elution Buffer (50 mM NaHCO_3_ and 1% SDS). After elution, ChIP sample was incubated with 10mg/mL proteinase K 4 h at 55 °C. Then DNA was purified using AMPure XP Beads (Beckman). Streptavidin C1 beads were used to capture biotinylated DNA. QIAseq FX DNA Library Kits were used to generate the sequencing library and HiChIP libraries were size-selected to 300–700 bp using AMPure XP beads (Beckman) and subjected to 2 × 50-bp paired-end sequencing on Hiseq XTen (Illumina).

### 4.3 ChIP-seq data processing and super enhancer analysis

Mapping of ChIP–seq data to hg19 was performed using bowtie2[38]. H3K27ac peak calling was performed using MACS1.4 with default parameter[39]. Peak calling for each sample was performed separately. Peaks that could not be identified in each colorectal cancer sample and peaks appeared within the region surrounding ± 2.5 kb of transcriptional start sites were excluded from any further analysis. Afterwards, the H3K27ac peaks of the 7 individual samples were merged into a single set of peaks. Super enhancers were identified using the rank ordering of super enhancers (ROSE) algorithm[14] In the case of colorectal cancer-specific enhancers, all peak regions were removed that contained any overlap with a peak detected in each normal colon region.

### 4.4 Unsupervised hierarchical clustering analysis

A matrix of the normalized H3K27ac density was generated using HOMER[40]. Variant enhancer loci (VELs) were defined as enhancers, which exhibited the greatest median absolute deviation (MAD) across all samples used for clustering. In the case of unsupervised hierarchical clustering between colorectal cancer and Roadmap Epigenomics samples data, 11,463 VELs were retained. These enhancers were used for unsupervised hierarchical clustering using a Pearson correlation as a distance metric.

### 4.5 *t*-SNE analysis

For clustering of H3K27ac ChIP–seq data from the colorectal cancer and normal colon cohorts together, we generated both cohorts normalized H3K27ac density matrix. Distance between samples was calculated by using 1-Spearman correlation coefficient as the distance measure. The resulting distance matrix was used to perform the *t*-SNE analysis (Rtsne package)

### 4.6 HiChIP data processing

Valid fragment pairs and interaction matrix was generated using the HiC-Pro[41]. Then the interaction loop calling was performed using the hichipper[42] and interaction loops in Figure 2B-E were shown as circos plot [43]. In the case of colorectal cancer specific enhancer loops detecting, the loops were picked that contained anchors overlapped with cancer specific enhancers.

### 4.7 HiC data analysis

Two HiC data sets in GSE133928, BRD3179 and BRD3179N [11], representing colorectal cancer tissue and normal colon tissue respectively, was analyzed to detect cancer and normal TADs. For the unchanged-boundary cancer TADs, the cancer TADs boundary was extremely overlapped with normal TADs. For the expand cancer TADs, the cancer TAD region contained only one complete normal TAD. For fusion cancer TADs, each cancer TAD region contained more than one complete normal TADs. For narrow cancer TADs, each cancer TAD was included in one normal TAD. For the split cancer TADs, more than one cancer TADs were included in one normal TADs. For the appear or disappear cancer TAD, the TADs existed only in cancer sample or normal sample respectively.

## Acknowledgments

This work was supported by the National Key R&D Program of China (NO.2017YFA0102600) and the Fundamental Research Funds for the Central Universities, Nankai University (63201087).

## Authors Contributions

B.C.,L.Z, conception and design, drafting or revising the article; B.C., L.Z., acquisition of data, analysis and interpretation of data; B.C., Y.M., A.H., J.B., W.W., and G.S. acquisition of data; B. C., analysis of all High-through sequencing data; B. C., W.W., acquisition of data and production of all bioinformatics figures; J.S, Z.Z, L.Z., revising the article.

## Availability of supporting data

Published data used in this study

The following published datasets were used in our analysis: GSE36204 for H3K27ac ChIP-seq analyses[15], GSE133928 for Hi-C data in colorectal cancer cells and normal colon cells [11] and RNA-seq data was gained from TCGA and GEPIA database[44].

## Data Availability

HiChIP data are deposited in the NCBI Gene Expression Omnibus (GEO, https://www.ncbi.nlm.nih.gov/geo/) under the accession number GSE173699.

